# Functional and sensitivity profiling of the *KIT* Mutation Landscape in Melanoma

**DOI:** 10.64898/2026.02.18.706482

**Authors:** Sai Fung Yeung, Mandy Sze Man Chan, Cherie Tsz Yiu Law, Alex Chun Ho Law, Carol Lee, Allen Ming Fai Leung, Mary Po Ki Chau, Helen Hoi Yin Chan, July Xi Chen, Ben C.B. Ko, Karen K.L. Chan, William C Cho, Stephen Kwok Wing Tsui

## Abstract

Melanoma in Asia presents a unique epidemiological profile, with a higher prevalence of acral and mucosal subtypes compared to Western populations. While *KIT* mutations are found in up to 15% of Asian melanoma cases, clinical outcomes with KIT inhibitors have been modest due to heterogeneous mutation profiles and a lack of specific patient selection criteria. This study characterizes the landscape of *KIT* mutations in melanoma using the GENIE database, identifying 86 recurrent hotspots, many of which are variants of unknown significance (VUS). We validated drug sensitivities for key mutations using *in vitro* and *in vivo* models. Our results indicate that while the L576P mutation is highly sensitive to multiple inhibitors, the N822K mutation shows resistance to imatinib but responds to sunitinib, nilotinib, and nintedanib. These findings highlight the necessity of genotype-guided therapeutic strategies and provide a rationale for future clinical trials combining broad-spectrum KIT inhibitors with immune checkpoint inhibitors.

**Translational Significance:** Melanoma subtypes prevalent in Asia, specifically acral and mucosal melanoma, frequently harbor *KIT* mutations but show poor response rates (23-26%) to the standard-of-care inhibitor, imatinib. This study challenges the current clinical practice of treating all *KIT*-mutated melanomas uniformly. We demonstrate that specific recurrent mutations, such as N822K, are intrinsically resistant to imatinib but highly sensitive to broad-spectrum inhibitors like sunitinib and nintedanib. By establishing a comprehensive “lookup table” of drug sensitivities for both common and previously uncharacterized *KIT* variants, this work provides the evidence base required to transition from a “one-size-fits-all” approach to a genotype-guided precision medicine strategy. Furthermore, validating these targets informs the design of next-generation clinical trials, particularly those combining optimal KIT inhibitors with immune checkpoint blockade to improve survival in currently underserved patient populations.

## Introduction

Melanoma in Asia have a significant higher mortality-to-incidence ratio compared to the Western Countries (IARC), with acral (AM) and mucosal melanoma (MM) subtypes being the most prevalent in Asia, accounting for 40-65% and 20-30% of cases, respectively (Cormier et al., 2006, Jang et al., 2014, Lee et al., 2012, Lv et al., 2016, Wang et al., 2016, Teh et al., 2018). Due to their rarity in the Caucasian population, there exist no specific treatment guidelines for advanced MM & AM currently, and they have shown less response to existing immunotherapy or other systemic treatments designated for cutaneous melanoma (Shoushtari et al., 2016, D’Angelo et al., 2017, Mao et al., 2021). Therefore, alternative treatment modalities and combination therapies are urgently needed to improve outcomes for Asian melanoma patients.

KIT mutations can be found in up to 15% of Asian melanoma cohorts (Kong et al., 2011, Sun et al., 2023, Jin et al., 2013), making it an interesting target for small molecule inhibitor therapy. Differ from GIST, the heterogeneity of mutation profiles and short-lived response toward KIT inhibitors as monotherapy in melanoma raise concerns (Janku et al., 2022). Previous experiences show tumours with KIT exon 11/13 mutations tend to be more sensitive to KIT inhibitors (Steeb et al., 2021), and broader-spectrum inhibitors may enhance treatment effectiveness in the diverse mutation landscape of melanoma (Janku et al., 2022, Kim et al., 2023).

Recent studies have shown intriguing benefits in combining anti-angiogenesis therapy with immune checkpoint inhibitors (ICI) for patients with Acral Melanoma (AM) and Mucosal Melanoma (MM). This combination yielded a median objective response rate (ORR) of 48% (Mao et al., 2022, Wang et al., 2023, Siming et al., 2022). Additionally, a tri-combination of camrelizumab, apatinib, and temozolomide resulted in a 64.0% ORR and significantly extended survival in advanced AM (Mao et al., 2023). Another unpublished study demonstrated remarkable ORR up to 90% in subgroups of patients (N=10) with KIT mutations in exon 11 when treated with a combination of imatinib and toripalimab (Si et al., 2022). Considering that most KIT inhibitors simultaneously target KIT and other angiogenic kinases such as VEGFR and PDGFR, we are interested in exploring the efficacy of combining broad-spectrum KIT inhibitors with ICI treatment in advanced melanoma patients with KIT mutations in the upcoming trial. This involves considering the sensitivity of KIT mutants to specific inhibitors and their anti-angiogenesis activity. Understanding the sensitivity of KIT mutants to inhibitors is essential as to minimize potential intrinsic KIT inhibitor resistance in tumors with certain specific KIT mutations (Janku et al., 2022). To facilitate this selection, we will also establish a high-throughput platform to catalogue all clinically recurrent KIT mutations and evaluate their sensitivity to various KIT inhibitors.

## Material and Methods

### Bioinformatics Analysis and Variant Selection

**AACR Project GENIE database (Version 12.0)** were queried to extract genomic profiles of over 130,000 tumor samples. *KIT* alterations, including point mutations, small insertions/deletions (indels), and structural variants, were filtered specifically for melanoma histology. Recurrent hotspots were defined as variants appearing in at least 3 independent cases. Variants were cross-referenced with the **ClinVar** and **COSMIC** databases to identify Variants of Unknown Significance (VUS). Structural modeling of mutant proteins was performed using existing crystal structures of the KIT kinase domain (PBD: 1T46) to predict the impact of mutations on the ATP-binding pocket and the activation loop.

### Cell Lines and Reagents

The IL-3-dependent murine pro-B cell line **Ba/F3** and the human melanoma cell line **MeWo** (ATCC) were used for functional characterization. Ba/F3 cells were cultured in RPMI 1640 supplemented with 10% FBS and 10% WEHI-3B conditioned medium (as a source of IL-3). MeWo cells were maintained in EMEM with 10% FBS. All cell lines were authenticated by STR profiling and confirmed mycoplasma-free via PCR-based testing. Small molecule inhibitors (Imatinib, Sunitinib, Nilotinib, Nintedanib, and Ripretinib) were purchased from MedChemExpress, dissolved in DMSO to a 10 mM stock, and stored at −80°C.

### Generation of Barcoded KIT Mutant Libraries

To achieve scalable functional testing, we utilized a **high-throughput mutation screening platform (HMPS)**. (i) **Vector Construction:** The wild-type *KIT* cDNA was cloned into the doxycycline-inducible pCW-Cas9-Blast lentiviral backbone (modified to remove Cas9). (ii) **Site-Directed Mutagenesis:** 86 recurrent *KIT* variants were generated using overlapping extension PCR. (iv) **Barcoding:** Unique 6-base pair (bp) DNA barcodes were inserted into the 3’ UTR of each mutant construct to allow for pooled competitive assays and precise quantification via next-generation sequencing (NGS). (iv) **Lentiviral Transduction:** Lentivirus was produced in HEK293T cells using psPAX2 and pMD2.G packaging plasmids. Target cells (Ba/F3 and MeWo) were transduced at a low multiplicity of infection (MOI < 0.3) to ensure single-copy integration and selected with blasticidin (10 mg/mL).

### Oncogenic Transformation and Cell Viability Assays

#### IL-3 Independence Assay

Ba/F3 cells expressing *KIT* mutants were washed three times in PBS to remove IL-3 and seeded in 96-well plates. Growth was monitored over 7 days; survival in the absence of IL-3 served as evidence of the mutant’s oncogenic transforming potential.

#### Drug Sensitivity (IC50)

Cells were treated with a 10-point serial dilution of KIT inhibitors. After 72 hours of incubation, cell viability was measured using the **CellTiter-Glo® Luminescent Cell Viability Assay** (Promega).

Luminescence was recorded using a microplate reader, and IC50 values were calculated using a non-linear regression model (log[inhibitor] vs. response) in GraphPad Prism 9.0.

### Western Blotting and Signaling Analysis

MeWo cells were induced with doxycycline (1 mg/mL) for 24 hours followed by 4 hours of serum starvation. Cells were then treated with indicated concentrations of inhibitors. Total protein was extracted using RIPA buffer supplemented with protease and phosphatase inhibitor cocktails. Equal amounts of protein (20–30 $\mu$g) were resolved by SDS-PAGE and transferred to PVDF membranes. Primary antibodies against **p-KIT (Tyr719), total KIT, p-ERK1/2, total ERK, p-AKT (Ser473), total AKT, p-STAT3, and p-SRC** (Cell Signaling Technology) were used for detection. beta-actin served as the loading control.

### In Vivo Xenograft Studies

All animal procedures were approved by the Institutional Animal Care and Experimentation unit.

#### Tumor Inculation

5 × 10^6 MeWo cells (expressing different KIT mutants) were resuspended in a 1:1 mixture of PBS and Matrigel and injected subcutaneously into the dorsal flank of 6-week-old female BALB/c nude mice.

#### Treatment Regimen

Once tumors reached an average volume of 25–30 mm^3^, mice were randomized (n=3 per group) to receive vehicle (0.5% methylcellulose), Imatinib (50 mg/kg), Sunitinib (40 mg/kg), or Nintedanib (50 mg/kg) via daily oral gavage.

#### Monitoring

Tumor volume (V = length x width^2^ / 2) and body weight were measured every 2 days. At the study endpoint, mice were euthanized, and tumors were harvested for IHC and molecular analysis.

### Statistical Analysis

Data are presented as mean +/- SEM. Differences between groups were analyzed using one-way or two-way ANOVA followed by Tukey’s post-hoc test for multiple comparisons. P < 0.05 was considered statistically significant. All analyses were performed using GraphPad Prism.

## Results

### Clinical Validation of KIT Inhibitor Efficacy in Exceptional Responders

To establish the clinical relevance of targeting specific *KIT* mutations in melanoma, we performed *in vivo* testing of sunitinib using xenograft models expressing KIT p.W557G and p.L576P variants in MeWo cells. KIT mutatns significantly accelerated tumor growth compared to EGFP controls and wild-type (WT) KIT (Fig. 1A-B). In these models, sunitinib treatment led to robust tumor regression.

**Figure 1.**
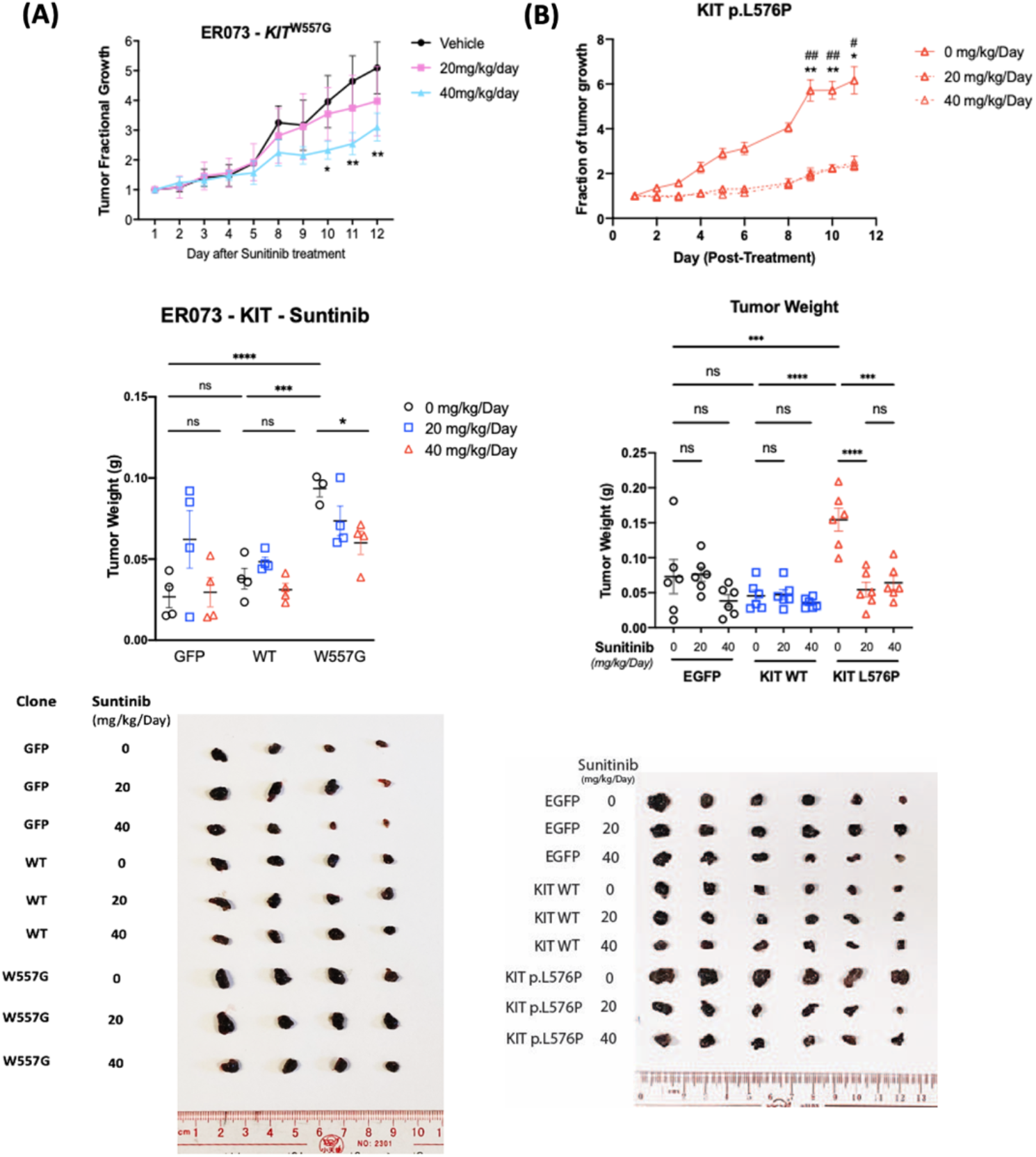
*In vivo* confirmation of Sunitinib efficacy against MeWo xenograft expressing wild-type (WT) KIT and/or KIT mutants derived from exceptional responders. (A) Tumor response of MeWo-EGFP, KIT-WT, and KIT p.W557G modeling exceptional responder no. 73 (N=2). (B) Tumor response of MeWo-EGFP, KIT-WT, and KIT p.L576P.

### The Genomic Landscape of KIT Mutations in Melanoma

Analysis of the GENIE v12 database was used to map the distribution of *KIT* alterations across over 130,000 tumor samples. We identified that *KIT* mutations in melanoma cluster predominantly within the juxtamembrane domain (Exon 11) and the kinase domain (Exons 13, 17, and 18) (Fig. 2A). A comparative pan-cancer analysis revealed that while some mutations overlap with Gastrointestinal Stromal Tumors (GIST), melanoma possesses a distinct enrichment of specific variants, such as those in the juxtamembrane region, which may dictate unique therapeutic sensitivities compared to other KIT-driven malignancies (Fig. 2B).

**Figure 2.**
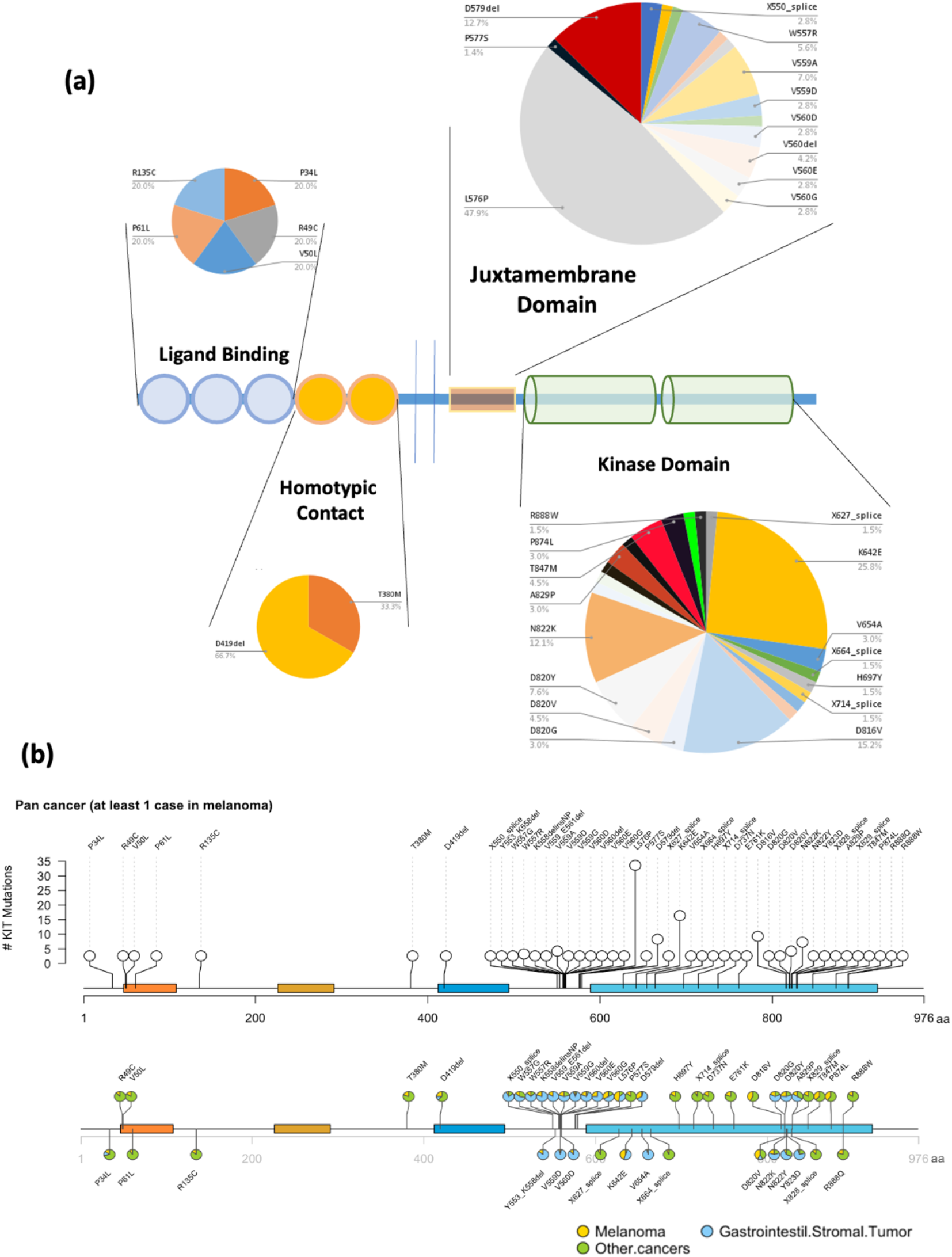
Mutation spectrum of KIT obtained from GENIE v12. (A) Schematic diagram depicting the distribution of recurrent KIT mutations identified in melanoma across the pan-cancer spectrum. The pie chart illustrates the relative abundance of mutations in specific domains of the KIT gene in melanoma. (B) Lolliplot plot presenting the relative abundance of specific mutations in melanoma (yellow), GIST (blue), and other cancer types in the pan-cancer dataset, as depicted in the pie chart.

### KIT Variants Drive Oncogenic Transformation and Constitutive Signaling

We next characterized the functional impact of these recurrent mutations. Western blot analysis in MeWo cells confirmed that the expression of *KIT* mutants leads to the constitutive phosphorylation of KIT (Tyr719) and the activation of downstream pro-growth and survival pathways, including ERK1/2, AKT, STAT3, and SRC (Fig. 3A). In 3D culture, MeWo cells harboring *KIT* mutations formed significantly larger and more invasive spheroids compared to WT controls over an 8-day period (Fig. 3B). Furthermore, when expressed in IL-3-dependent Ba/F3 cells, these *KIT* variants conferred IL-3 independence, confirming their status as potent oncogenic drivers (Fig. 3C), indicative for its oncogenic properties.

**Figure 3.**
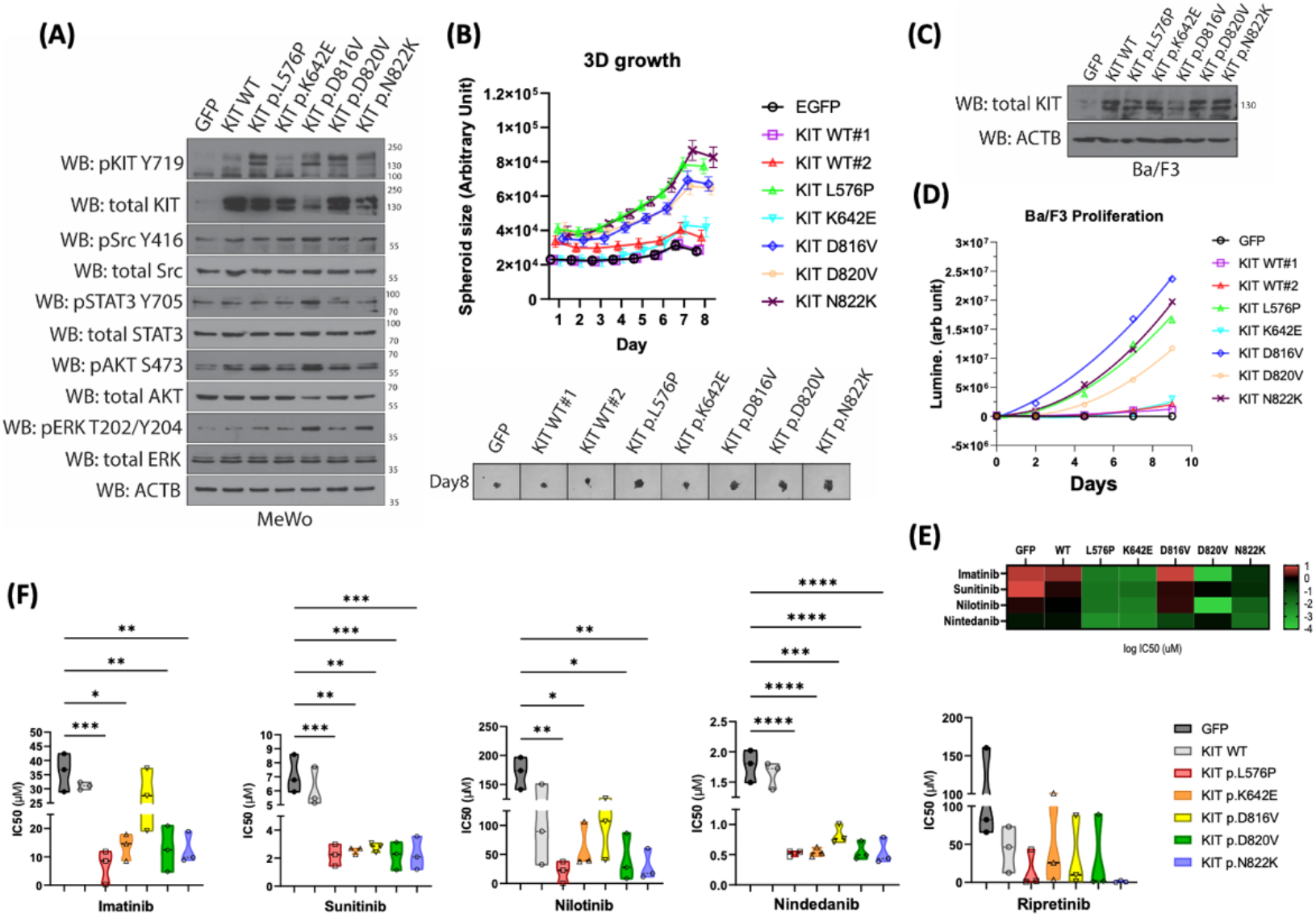
KIT mutations shows differential responses towards common KIT inhibitors in vitro. (A) Western Blot analysis revealed changes in cell growth and proliferation pathways upon expression of KIT mutants in MeWO melanoma cells. (B) Upper panel: Monitoring relative growth of melanoma cell spheroids expressing KIT mutants using high-content microscopy. Lower panel: Representative image depicting spheroid size after 8 days of growth in 3D environment. (C) Western Blot demonstrating the expression of KIT in Ba/F3 cells. (D) Assessment of relative growth of Ba/F3 cells expressing different KIT mutants following withdrawal of IL-3 from the culture medium, measured using cell-titre glo assay. (E) Heatmap shows the relative IC50 of Ba/F3 cell with KIT mutants against KIT inhibitors. (F) Violin plot illustrating the IC50 values of MeWo cells expressing KIT mutants when treated with various KIT inhibitors. Results were obtained from three independent experiments. The horizontal line represents the mean, and statistical analysis was performed using one-way ANOVA.

Next, We conducted comprehensive drug sensitivity profiling across a panel of clinical-grade KIT inhibitors. Heatmap showing the IC50 values in Ba/F3 models and violin plots in MeWo models revealed a high degree of heterogeneity in drug response (Fig. 3D-F). KIT p.L576P exhibited broad sensitivity to most inhibitors tested, including imatinib and sunitinib. In contrast, KIT p.N822K showed a “selective” sensitivity profile; while it was significantly less responsive to the frontline inhibitor imatinib, it remained highly vulnerable to second-generation inhibitors such as sunitinib, nilotinib, and nintedanib.

### In Vivo Anti-Tumor Efficacy and Resistance Patterns

The differential sensitivity observed *in vitro* was validated in mouse xenograft models. MeWo-KIT p.L576P tumors showed significant growth inhibition when treated with imatinib, sunitinib, nilotinib, and nintedanib (Fig. 4A-B). Crucially, the p.N822K mutant—a common variant in mucosal melanoma—demonstrated marked resistance to imatinib but was successfully suppressed by sunitinib and nintedanib (Fig. 4C). Additionally, we evaluated the “switch-control” inhibitor ripretinib, finding that its efficacy varied significantly depending on the specific *KIT* genotype, further emphasizing the need for mutation-specific treatment selection (Fig. 4D-F).

**Figure 4.**
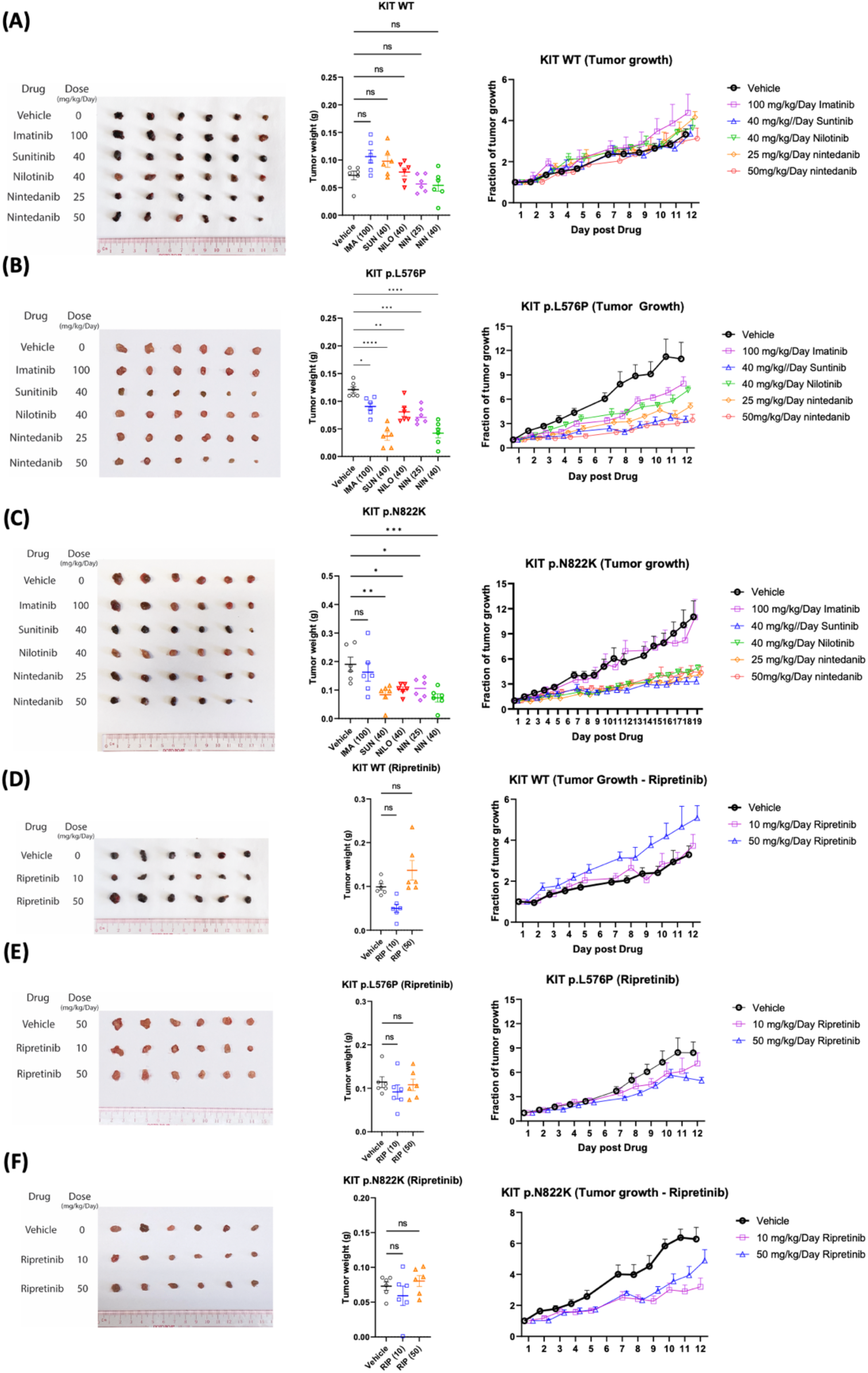
In vivo anti-tumor efficacy of KIT inhibitors against MeWo cells expressing KIT WT/mutants in a mouse xenograft model. (A-C) Response of MeWO-KIT-WT, MeWo-KIT p.L576P, and MeWo-KIT-p.N822K mouse xenograft models to Imatinib, Sunitinib, Nilotinib, and Nintedanib, respectively (n = 3 independent experiments). (D-F) Response of MeWO-KIT-WT, MeWo-KIT p.L576P, and MeWo-KIT-p.N822K mouse xenograft models to Ripretinib (n = 3 independent experiments). Fraction of tumor growth represents the change in tumor volume normalized to day 0 of treatment.

### Generation of a High-Throughput Mutation Screening Platform (HMPS)

To systematically address the 86 recurrent *KIT* variants (including 48 VUS) identified in our genomic analysis, we developed a scalable High-Throughput Mutation Screening Platform (HMPS) (Fig. 5). This workflow utilizes universal primers and a three-step PCR reaction to generate a lentiviral library of *KIT* mutants, each tagged with a unique 6-bp barcode. Following low-MOI transduction into experimental cell lines, this system allows for the simultaneous evaluation of the transforming potential and drug sensitivity of dozens of variants in a single pooled experiment using barcoded DNA sequencing in the future.

**Figure 5.**
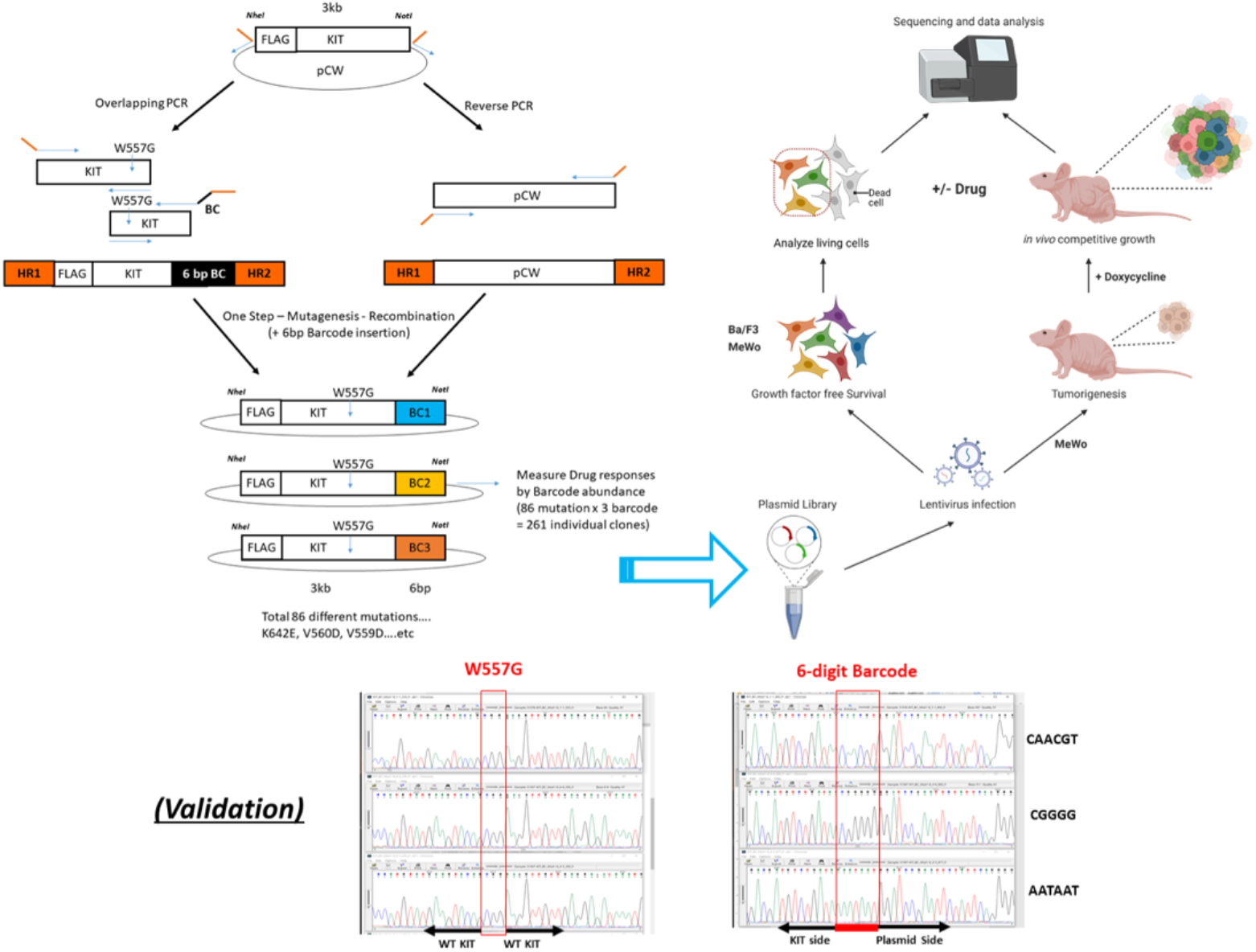
Schematic representation of detailed workflow for high-throughput mutation screening platform. Left side: A scalable and standardized workflow was established to generate a lentiviral KIT mutant library containing barcode sequences with 6 base pairs. Mutated ORFs were generated using universal primers and a three-step PCR reaction, followed by renomination-mediated cloning into an inducible lentiviral vector. Right side: Transformed experimental cells were evaluated for their transformation potential or drug sensitivity under induction with puromycin. The successful generation of mutant variants was confirmed through genomic DNA sequencing.

## Discussion

The off-label use of imatinib as the default therapy for *KIT*-mutated melanoma appears to be a carryover from its success in GIST. However, our comparative analysis highlights a critical divergence: while GIST mutations are heavily clustered in exon 11, melanoma exhibits a much higher frequency of kinase domain mutations, such as N822K. This distinction is not merely academic; it has profound pharmacological consequences. Our data reveals that while juxtamembrane mutations like L576P retain broad sensitivity, activation-loop mutations like N822K are intrinsically resistant to imatinib. This structural and functional heterogeneity provides a clear biological explanation for why unstratified phase II trials have historically struggled. The discovery that nintedanib and sunitinib can effectively “bypass” this resistance by suppressing N822K-driven growth *in vivo* offers an immediate and actionable alternative to the current, failing clinical paradigm.

Interestingly, sunitinib and nintedanib often outperformed more selective inhibitors, even against imatinib-sensitive variants. This suggests that the therapeutic “sweet spot” in melanoma may not lie in absolute selectivity. Given the highly vascular nature of mucosal and acral subtypes, the efficacy of these agents likely stems from a dual-threat mechanism: direct blockade of the oncogenic KIT axis (p-STAT3/p-ERK) coupled with the disruption of the tumor microenvironment through VEGFR and PDGFR inhibition. This multifaceted assault may be exactly what is needed to delay the rapid onset of resistance that plagues single-target TKI therapy.

One of the most significant hurdles in precision oncology is the clinical “limbo” of Variants of Unknown Significance (VUS). Our analysis shows that over half of recurrent *KIT* mutations in melanoma fall into this category, leaving clinicians with no evidence-based treatment path. The development of our High-Throughput Mutation Screening Platform (HMPS) is an attempt to resolve this ambiguity at scale. By moving from anecdotal case reports to a standardized, barcoded library of all 86 recurrent variants, we can systematically map the drug sensitivity of the entire *KIT* landscape. This work transforms a collection of genomic data into a functional “lookup table,” providing the empirical evidence needed to guide drug selection in real-time.

The data presented here argue for an urgent pivot in clinical trial design. The era of single-arm imatinib trials for any *KIT* mutation should be over. Instead, we propose a transition to “matching” or “umbrella” trials where patients are stratified by their specific genotype such as

1. **Imatinib-sensitive hotspots** (e.g., L576P) receive potent, potentially multi-targeted KIT inhibitors.
2. **Imatinib-resistant variants** (e.g., N822K, D816V) are immediately directed to agents like nintedanib or ripretinib.
3. **Combination strategies:** Perhaps most importantly, these matched TKIs should be investigated as backbones for combination with immune checkpoint inhibitors. The synergy between TKI-induced antigen release and PD-1 blockade may finally provide the durable responses that have been so elusive in this patient population.

Ultimately, this study is expected to serve as a functional and pharmacological roadmap, transitioning away from a universal treatment model toward a more precision strategies outlined here is the most viable path to improving survival for patients with acral and mucosal melanoma.

